# Arsenite methyltransferase 3 regulates hepatic energy metabolism which dictates the hepatic response to arsenic exposure

**DOI:** 10.1101/2023.04.05.535637

**Authors:** Patrice Delaney, Nouf Khan, Matthew J. O’Connor, Elizabeth Mayela Ambrosio, Anna Garcia-Sabaté, Jeremy C. M. Teo, Spiros A. Pergantis, Elke Ober, Kirsten C. Sadler

**Affiliations:** Biology Program, New York University Abu Dhabi, Saadiyat Campus, P.O. Box 129188, Abu Dhabi, United Arab Emirates; Core Technology Platform, New York University Abu Dhabi, Saadiyat Campus, P.O. Box 129188, Abu Dhabi, United Arab Emirates; Engineering Division, New York University Abu Dhabi, Saadiyat Campus, P.O. Box 129188, Abu Dhabi, United Arab Emirates; Department of Chemistry, University of Crete, Heraklion, Crete, 70013, Greece; Danish Stem Cell Center (DanStem), University of Copenhagen, Copenhagen, Denmark; Center for Genomics and Systems Biology (CGSB), New York University Abu Dhabi, P.O. Box 129188, Abu Dhabi, UAE

**Author notes:** To whom correspondence should be addressed: Kirsten C. Sadler, New York University Abu Dhabi, P.O. Box 129188, Abu Dhabi, United Arab Emirates.

**Keywords:** Arsenite Methyltransferase (As3mt), Zebrafish, Inorganic Arsenic, Liver, Hepatoxicity, Fatty Liver

## Abstract

Inorganic arsenic (iAs(III)) is among the most pervasive environmental toxicants in the world. The iAs metabolizing enzyme, arsenite methyltransferase (*AS3MT)*, is a key mediator of iAs(III) toxicity and has been almost exclusively investigated in the context of iAs(III) exposure. We use functional genomics approach with zebrafish *as3mt* mutants which lack arsenite methyltransferase activity to uncover novel, arsenic-independent functions for As3mt. Transcriptomic analysis of untreated whole larvae, and the larval and adult livers from *as3mt* mutants revealed thousands of differentially expressed genes (DEGs) compared to wild-type controls. These were enriched for genes functioning in the ribosome or mitochondria. Nearly all genes in the citric acid cycle and mitochondrial transport were downregulated in *as3mt* mutant livers. This resulted in reduction in reactive oxygen species levels by half and fatty liver in 81% of *as3mt* mutant larvae. An inverse expression pattern was detected for over 2,000 of the As3mt regulated DEGs in the liver of larvae with transgenic overexpression of As3mt in hepatocytes. Replacing *as3mt* expression in hepatocytes of *as3mt* mutants prevented fatty liver, demonstrating that As3mt has novel, cell-autonomous and arsenic-independent functions regulating mitochondrial metabolism. We suggest that these functions contribute to iAs toxicity, as the mitochondrial function genes that were downregulated in the liver of unexposed *as3mt* mutants were further downregulated upon iAs exposure and *as3mt* mutants were sensitized to iAs. This indicates that As3mt regulates hepatic energy metabolism and demonstrates that, in addition to its role in iAs detoxification, the physiological functions of As3mt contribute to arsenic toxicity.

**SIGNIFICANCE:** Arsenic is an endemic environmental toxicant, and the current paradigm is that susceptibility to arsenic toxicity is dictated by levels of expression of the arsenite 3 methyltransferase gene (As3mt), which is dedicated enzyme involved in arsenic detoxification. Our data showing that As3mt serves arsenic-independent functions in energy metabolism challenge this paradigm. We show that zebrafish *as3mt* mutants have loss of mitochondrial function and develop fatty liver and suggest that *as3mt* mutants are sensitized to arsenic toxicity due, in part, to impaired mitochondrial function. This finding opens an entirely new area of study to identify the cellular function of As3mt and further advances the understanding of how genetic variants in As3mt confer sensitivity arsenic toxicology.

## INTRODUCTION

The World Health Organization (WHO) has placed iAs(III) within the top ten chemicals of major public concern because roughly a third of the world’s population is at risk of exposure to this devastating toxicant (1). Exposure to iAs is associated with major health ramifications, including multiple forms of cancer, skin lesions, cardiovascular disease, neurological defects, diabetes and liver disease (2-6). Arsenic metabolizing genes are highly conserved in metazoa (7); in humans and other vertebrates, arsensite methyltransferase (AS3MT) catalyzes trivalent iAs to mono- or dimethylated arsenic (MMA, DMA) using s-adenosyl methionine (SAM) as a donor and glutathione (GSH) as a reducing agent. This is hypothesized to reduce retention of iAs by converting ionic bonds into more stable covalent bonds, preventing promiscuous interactions with other cellular components and increasing the arsenic clearance rate (3). While As3mt activity can reduce these damaging effects of iAs(III), the process of iAs metabolism by As3mt can deplete SAM and cellular antioxidant stores, and generate reactive oxygen species (ROS), all of which have been proposed to contribute to arsenic toxicity in humans (8) and animal models (9-12).

The importance of As3mt in arsenic-induced health outcomes is predicted on the finding that single nucleotide polymorphisms (SNPs) in the *AS3MT* locus correlate with differential susceptibility to iAs toxicity in humans. For example, communities in the Andes Mountains, which have historically resided near rivers with arsenic concentrations 100 fold higher than the WHO’s safe limit of 10 µg/L, have *AS3MT* haplotypes associated with more efficient methylation of iAs and partial resistance to iAs (13). Many protective SNPs are found within *AS3MT* introns, and some SNPs increase expression of *AS3MT* which is proposed as the mechanism increasing iAs methylation capacity, decreasing iAs retention and lowering its toxic effects (13). In support of this, one study found an association between increased plasma *AS3MT* expression and putative protective *AS3MT* SNPs whereas another discovered correlation between reduced iAs metabolism efficacy and reduced *AS3MT* expression in people with *AS3MT* haplotypes associated with iAs susceptibility (14, 15). Studies of *As3mt* knock-out (KO) mice support this, as these have higher iAs retention and toxicity compared to wild-type (WT) mice (16).

The prevailing paradigm is that the sole function of AS3MT is to methylate arsenic. Several lines of evidence suggest an alternative: AS3MT is highly conserved, is expressed in a tissue specific fashion in the absence of iAs, and *As3mt* KO mice have metabolic changes (17, 18), with differences in the levels of specific phosphatidylcholine (PC) species (18) and cultured cells with *AS3MT* knockdown decreases proliferation (19). Moreover, other genome-wide association studies have found that variants of an AS3MT isoform correlates with schizophrenia (20, 21) and depletion of AS3MT in neurons resulted in the differential expression of over 1,400 genes (22). However, in the decades since AS3MT was discovered, aside from these few studies, its role in cellular homeostasis has not been investigated.

Here, we present a functional genomics study with *as3mt* mutant (*as3mt*^*-/-*^) zebrafish that uncovers an entirely new function for As3mt independent of its role in iAs metabolism. Transcriptomic analysis of *as3mt*^*-/-*^ whole larvae, and livers from larvae and adults revealed that over 3,000 genes were differentially expressed in the absence of arsenic. The downregulated genes were enriched for functions in the ribosome and mitochondrial membrane. Transgenic expression of *as3mt* in hepatocytes increases the expression of these same genes, including all those that function in the mitochondria. This indicates As3mt as a direct regulator of these genes and pathways in the liver. Importantly, these transcriptomic changes have functional consequences as we found lower levels of ROS produced by *as3mt*^*-/-*^ mutants, and nearly all develop steatosis by 120 hours post-fertilization (hpf). Steatosis was rescued by transgenic expression of *as3mt* only in hepatocytes, demonstrating a cell autonomous function of As3mt in hepatic lipid metabolism. Importantly, iAs is more toxic to *as3mt* mutant larvae compared to WTs. Despite a largely similar transcriptional response to iAs exposure, most of same mitochondrial genes that were downregulated in unexposed *as3mt*^*-/-*^ mutants are further decrease in expression following iAs exposure. This provides direct evidence of novel functions of As3mt in energy homeostasis and we proposed that, in addition to the essential role in iAs metabolism, the function of As3mt in cellular metabolism influence susceptibility to arsenic.

## RESULTS

### As3mt is highly expressed in hepatocytes and is enzymatically active in zebrafish larvae

Our previous studies in zebrafish demonstrated that *as3mt* mRNA is maternally provided and dynamically expressed during zebrafish development, with expression enriched in the liver by 120 hpf (11). We used a gene trap reporter line (23) for *as3mt (Tg(UAS:GFP;gSAlzGFFD886A))* which shows that expression is high and restricted to the liver as early as 72 hpf (**Figure S1A**-**1B**). RNAseq analysis of single livers dissected from WT 120 hpf larvae and adults shows that *as3mt* falls within the top ∼4% of genes expressed in the zebrafish liver (**Figure S1C**). Immunofluorescence of the WT liver at 120 hpf shows that As3mt is localized to punctate structures in hepatocytes (**Figure S1D)**.

We established As3mt activity in zebrafish larvae by treating embryos from 6-120 hpf with 1 mM sodium arsenite (iAs(III)) using a previously optimized protocol (12, 24). Ion chromatography (IC) inductively coupled plasma mass spectrometry (ICP-MS) was used to detect As(III), its oxidized form As(V), and the metabolic products DMA and MMA, with a detection range from 0.75 – 12.5 ppb of arsenic species. As expected, no arsenic species were detected in untreated larvae and DMA and MMA were detected in treated larvae, albeit at reduced levels compared to As(III) (**Figure S1E**). We also detected a tri-methylated arsenical species, arsenobetaine (AsB), which is abundant in marine animals and some freshwater fish, and is proposed to be generated by bacterial constituents of the microbiome (25). Together, these data show that in zebrafish larvae, as in mammals (26, 27), As3mt is specifically and highly expressed in the liver and is enzymatically active.

### Loss of *as3mt* increases iAs accumulation and toxicity in zebrafish larvae

We generated zebrafish *as3mt* mutants using CRISPR/Cas9 using an sgRNA targeting exon 3 of the *as3mt* locus (**Figure S2A**). We identified an allele with eight base pair indel which causes a frameshift that generates a novel 13 amino acid sequence following amino acid 3 and a premature stop codon predicted to truncate the protein upstream of the catalytic domain (**Figure S2B-F**). We bred these to homozygosity and, as predicted based on the lack of overt phenotypes in *As3mt* knock out mice (16, 28), *as3mt*^*-/-*^ mutants showed no significant difference in development, morphology, size, reproductive capacity or behavior (**Figure 1A-C** and not shown). Exposure to 1 mM iAs(III) from 6-120 hpf and assessment As(III), DMA, MMA and As(V) levels using IC-ICP-MS showed markedly increased levels of As(III) and undetectable levels of DMA and MMA in *as3mt*^*-/-*^ mutants (**Figure 1D**). This demonstrates that this is null or strongly hypomorphic allele.

**Figure 1.**
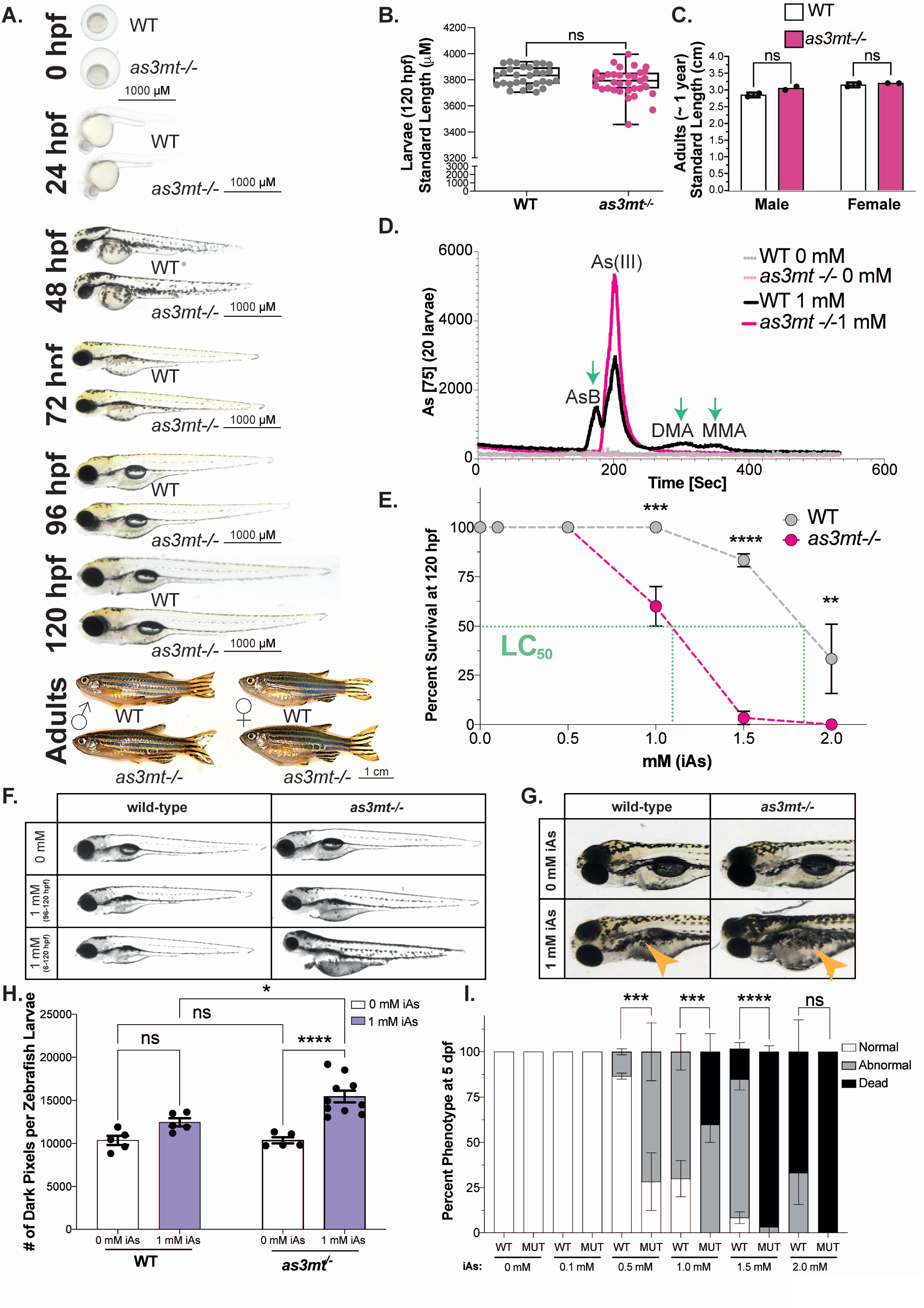
Loss of *as3mt* increases iAs accumulation and sensitivity in zebrafish larvae. **A**. Representative images of WT and *as3mt*^*-/-*^ larvae and adults. **B-C**. Standard length of larvae (μM; B) and adults (cm; C). **D**. Average IC-ICP-MS counts of AsB, AsIII, DMA, and MMA from WT (grey) or *as3mt*^*-/-*^ (pink) zebrafish extracts after 0 (dotted lines) or 1 mM iAs (6-120 hpf) challenge (solid lines) (n = 40, 2 clutches). **E**. Percent survival of WT (grey) and *as3mt*^*-/-*^ (pink) across 0, 0.5, 1.0, 1.5 and 2.0 mM iAs exposure from 6-120 hpf. Intersect with green line denotes the LC_50_. n = ∼60, 3 clutches. **F**. Representative images of WT and *as3mt*^*-/-*^ larvae during 0 mM, 1 mM acute (96-120 hpf), or 1 mM chronic (6-120 hpf) exposure to iAs. **G**. Images of 120 hpf zebrafish pigmentation at 0 mM or 1 mM chronic (6-120 hpf) iAs challenge (purple). Orange arrows mark pigmentation clusters. **H**. Quantification of pigmentation per larvae (n = 5-10, 1 clutch). **I**. Percent phenotype (normal (white), abnormal (grey) or dead (black) after 0-2 mM iAs exposure from 6-120 hpf (n = ∼60, 3 clutches).

*As3mt* KO mice are more sensitive to the toxic effects of iAs (16, 28) and we find that exposing *as3mt*^*-/-*^ mutants and WT embryos to a range of iAs(III) concentrations from 6-120 hpf reduced the lethal concertation 50 (LC_50_) from ∼1.9 mM iAs(III) in WT larvae to ∼1.1 mM in *as3mt*^*-/-*^ mutants (**Figure 1E**). At lower concentrations of iAs(III), *as3mt*^*-/-*^ mutants developed more severe edema, grey yolk **(Figure 1F**) and melanocyte expansion (**Figure 1G-H**) compared to WT embryos. These effects were dose dependent and significantly increased in mutants (**Figure 1I**). Thus, while loss of *as3mt* does not result in any gross morphological defects in zebrafish embryos, it significantly increases iAs(III) retention and toxicity.

### Thousands of genes deregulated in *as3mt* mutants

We hypothesized that if *as3mt* has physiological functions that are independent of iAs(III) metabolism then the loss of As3mt would induce a transcriptional response in the absence of iAs exposure. We performed RNAseq analysis on pools of WT and *as3mt*^*-/-*^ mutant whole larvae and livers at 120 hpf and normalized gene expression of mutants to WT controls (**Figure 2A**). This uncovered 3,400 differentially expressed genes (DEGs) in whole *as3mt*^*-/-*^ mutant larvae and 2,812 DEGs in mutant livers (padj < 0.05; **Figure 2B; Supplemental Table S1-2**), with 701 DEGs common to both datasets, 86% of which (608 genes) show a strong positive correlation in expression (r = 0.832; **Figure 2B** and **S3A**). To determine if this correlation extended to genes that were differentially expressed in one dataset but did not reach statistical significance in the other, we plotted the expression of the unified geneset of DEGs from both samples which showed a strong positive correlation (r = 0.640) of 60% of these genes (3,280 genes), with 1,457 genes that were upregulated and 1,823 genes downregulated in both samples (**Figure 2C)**. This indicates that the loss of As3mt disrupts cellular homeostasis during zebrafish development, resulting in a marked transcriptional response.

**Figure 2.**
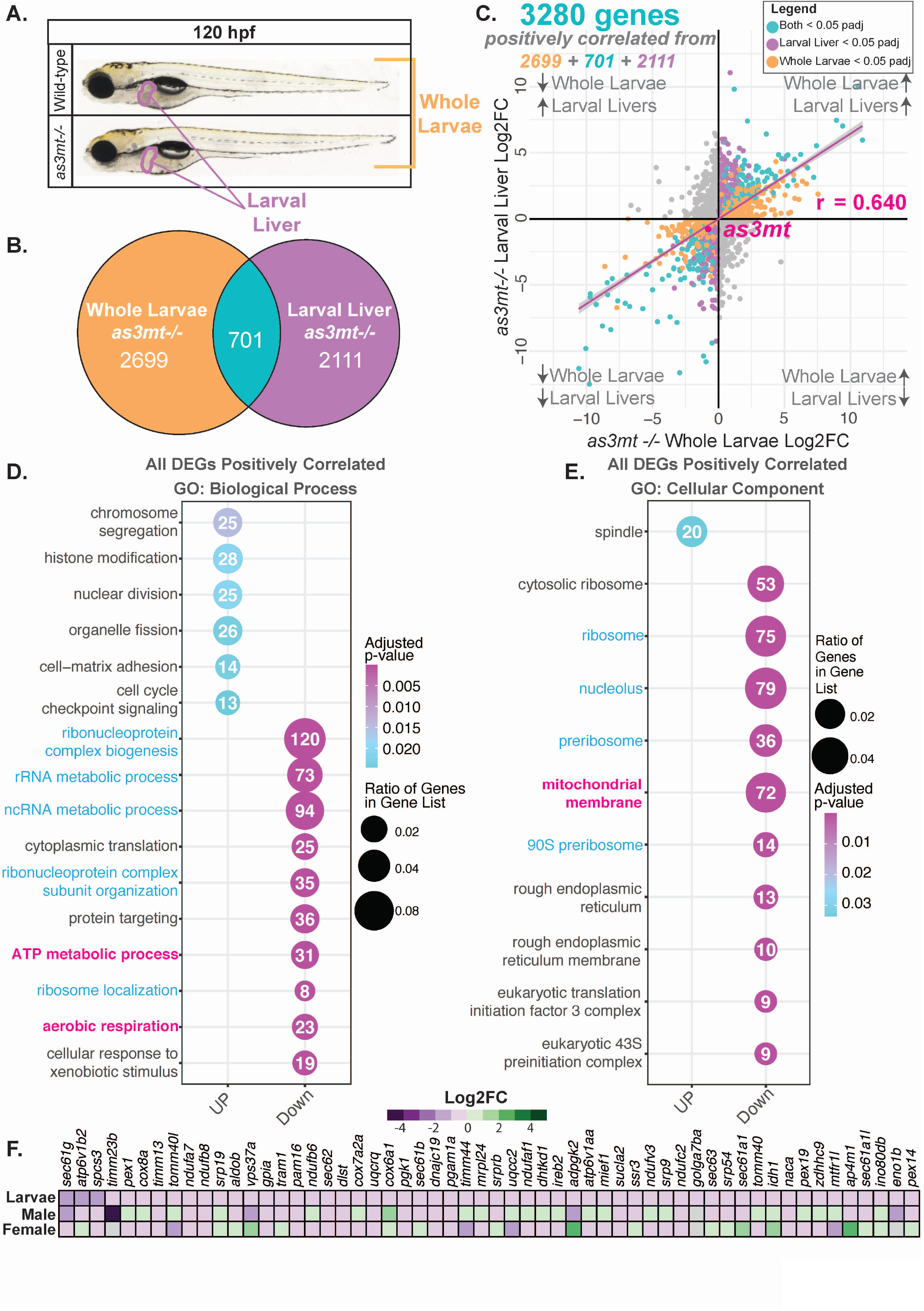
Physiological loss of *as3mt* elicits significant non-tissue-specific transcriptomic alterations at 120 hpf in zebrafish larvae. **A**. Representative images of 120 hpf WT and *as3mt*^*-/-*^ whole larvae (orange) and livers (outlined in purple). **B**. Overlap of differentially expressed genes (DEGs) from pooled *as3mt*^*-/-*^ whole larvae (orange) or larval livers (purple) RNAseq datasets. **C**. Cross plot of DEGs in WT and *as3mt*^*-/-*^ with negatively correlated genes in grey. Pearson’s correlation coefficient (r) and *as3mt* expression are marked in magenta. **D-E**. GO BP (D) and CC (E) of all positively correlated genes. **F**. Heatmap of Log2FC of genes from unified set from mitochondria genes in D (highlighted in pink) in larval, adult male and adult female livers.

Gene ontology (GO) analysis of these 3,280 positively correlated genes revealed that while the upregulated genes were enriched for a variety of cellular functions, nearly all of the downregulated genes were enriched in pathways related to ribosome biogenesis and function, ATP generation and xenobiotic metabolism (**Figure 2D**) and functioned in the in ribosome or mitochondrial membrane (**Figure 2E**). This same pattern was detected in the shared, positively correlated DEGs (**Figure S3B**). Interestingly, ribosome function and translation were also enriched in AS3MT deficient neurons (22). This suggests that loss of As3mt during development deregulates ribosome function, protein translation and mitochondrial function.

To determine if these changes persisted through adulthood, we performed RNAseq analysis on livers from male and female WT and *as3mt*^*-/-*^ mutant adult zebrafish. This showed sex specific differences in gene expression in both *as3mt*^*-/-*^ mutants and WTs (**Figure S4A-B**). The expression pattern in mutant males shows similar expression profile to larval livers, with 212 DEGs common to the *as3mt*^*-/-*^ mutant adult male livers, and nearly all positively correlated in their expression in both samples (**Figure S3C-D Table S3**). Importantly, all the downregulated DEGs in *as3mt*^*-/-*^ mutant larval livers that function in ATP synthesis and aerobic respiration were also downregulated in adult male *as3mt*^*-/-*^ livers (**Figure 2F**). Key genes regulating metabolic functions, including *vdac1*, which is essential for mitochondrial membrane transport (29), and *pemt*, which is required for PC synthesis (30) and for VLDL secretion (31), were significantly downregulated in larval and adult liver samples (**Figure S4F**). These data show that loss of *as3mt* elicits lifelong downregulation of genes that play important roles in mitochondrial and lipid metabolism in the liver, suggesting that these functions are perturbed by As3mt deficiency.

### Cell autonomous function of As3mt in hepatocyte energy metabolism

To address the hepatocyte specific role of *as3mt*, we generated a transgenic zebrafish line with moderate (1.5-2 fold) overexpression of zebrafish *as3mt* in hepatocytes ((*Tg(fabp10a:As3mt;cryaa:dsRed)* hereafter referred to as Tg-*as3mt*^*wt/tg*^; **Figure 3A-B**) which had no measurable effect on hepatic, biliary or endothelial cell morphology, hepatic architecture or liver function at 120 hpf (**Figure S5A-H**). RNAseq analysis of livers from 120 hpf Tg-*as3mt*^*wt/tg*^ larvae detected 139 DEGs (**Figure 3C**), 22 of which overlapped with the DEGs detected in *as3mt*^*-/-*^ mutant larval livers (**Figure 3D**). Analysis of the unified geneset of all DEGs from both *as3mt*^*-/-*^ mutant and Tg-*as3mt*^*wt/tg*^ livers (2,929 genes; **Table S4**) showed that 60% (1,780 genes) of the genes upregulated in mutants were downregulated in transgenics, and *vice versa* (r = -0.177) (**Figure 3E**). GO analysis of the negatively correlated genes revealed enrichment of ATP generation, respiration and metabolite precursor generation which were up in Tg-*as3mt*^*wt/tg*^ and down in *as3mt*^-/-^ livers (**Figure 4F**). UPSET analysis of the genes in these pathways demonstrated that many were shared between all the pathways involved in mitochondrial metabolism (**Figure 4G**). Notably, most genes functioning in the citric acid cycle (TCA) (**Table S5**) were downregulated in *as3mt*^*-/-*^ mutant livers and upregulated in Tg-*as3mt*^*wt/tg*^ livers (**Figure 3H**). We conclude that mitochondrial transport and ATP generation in zebrafish hepatocytes is directly and specifically regulated by As3mt.

**Figure 3.**
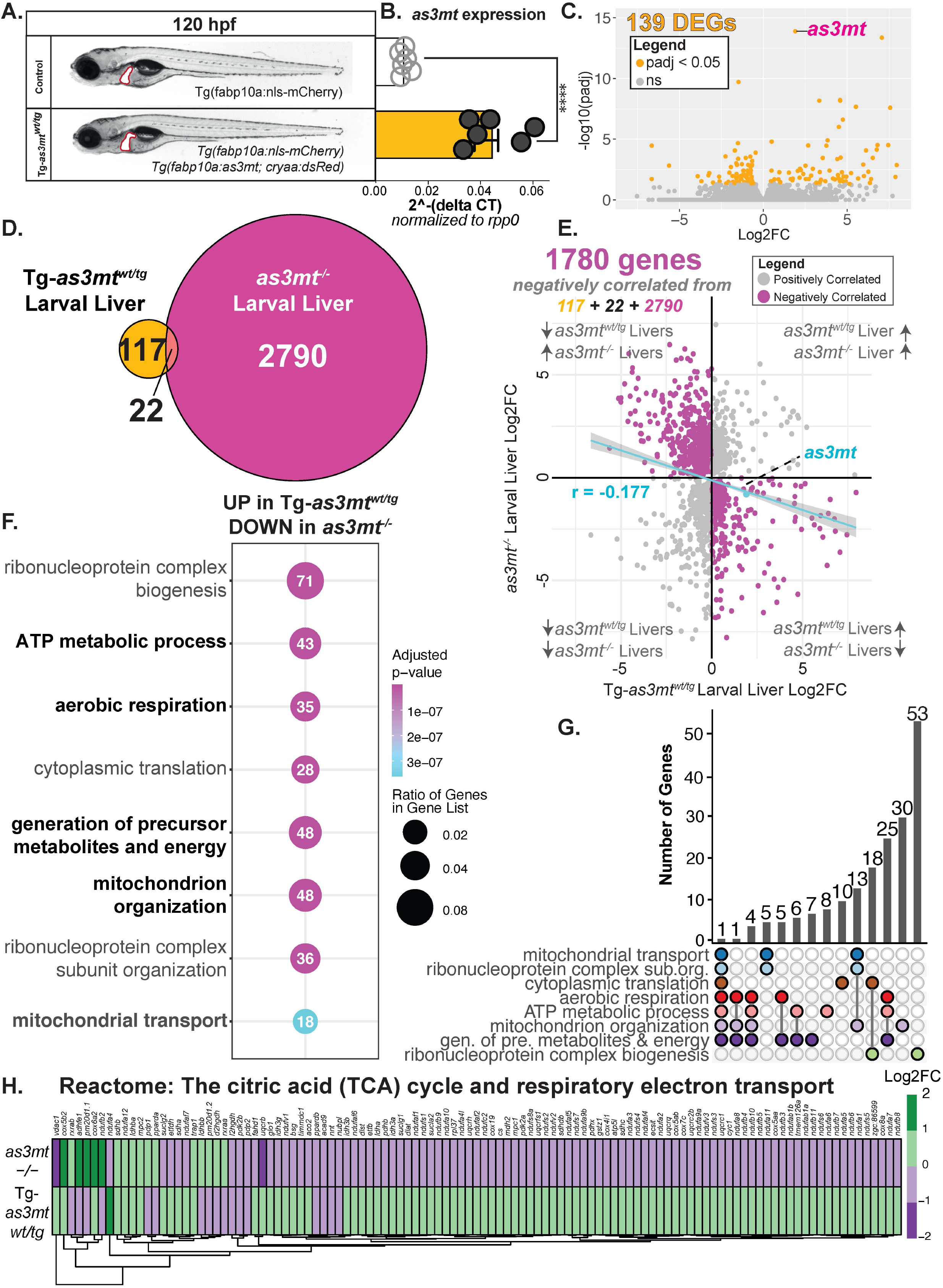
Loss or gain of *as3mt* is sufficient to alter TCA gene expression. **A**. Representative images of transgenic zebrafish without (top; control) and with *as3mt* hepatic overexpression (*tg:fabp10a:as3mt, cryaa:DsRed;* Tg-*as3mt*^*wt/tg*^) (bottom). **B**. qPCR analysis of singe livers micro-dissected from 120 hpf WT (white) or Tg-*as3mt*^*-wt/tg*^ (gold) zebrafish larvae for *as3mt* expression. **C**. DEGs from 120 hpf Tg-*as3mt* ^*wt/tg*^ livers compared to WT. **D**. Overlap of DEGs in *as3mt*-/- (pink) and Tg-*as3mt* ^*wt/tg*^ (gold). **E**. Cross plot of all DEGs. Negatively correlated genes are pink in blue (386/686). Pearson’s correlation coefficient (r) and *as3mt* expression are marked in magenta. **F**. GO BP analysis of negatively correlated genes from **G**. UPSET plot genes from D. **H**. Heatmap of TCA cycle genes in *as3mt*^*-/-*^ (top) and Tg-*as3mt*^*-wt/tg*^.

**Figure 4.**
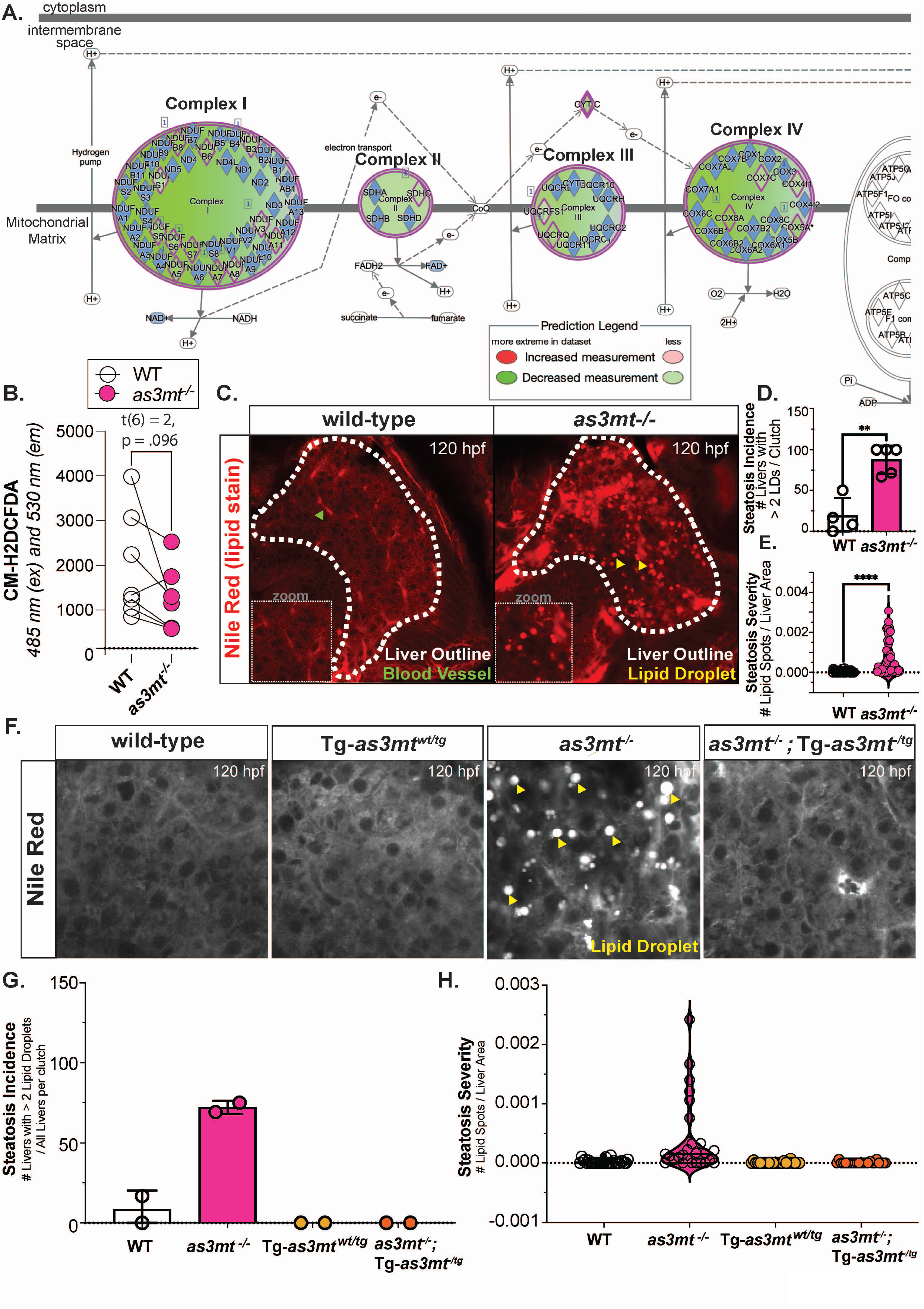
Loss of *as3mt* directly increases hepatic steatosis incidence and severity. **A**. IPA image of *as3mt*^*-/-*^ significant DEGs in oxidative phosphorylation pathway. **B**. ROS readouts from WT (white) or *as3mt*^*-/-*^ (pink) zebrafish larvae. **C**. Representative confocal images of WT (left) tand *as3mt*^*-/-*^ (right) larval livers stained with Nile red. **D**. Percent steatosis incidence from each clutch. **E**. Severity of steatosis (# of lipid droplets / liver area) in each liver. n = 35 livers per genotype, 4 clutches. pval = 0.0020, unpaired t-test. **F**. Representative images of livers in WT, *as3mt*^*wt/tg*^, *as3mt*^*-/-*^ and *as3mt*^*-/-;*^ Tg*-as3mt*^*wt/tg*^. Quantification of incidence (**G**) and severity (**H**) for each group.

### *as3mt* loss in hepatocytes reduces ROS and causes fatty liver

The gene expression profile shows expression of many genes that function in Complexes I-IV to be downregulated in *as3mt*^*-/-*^ mutant livers, and Ingenuity Pathway Analysis predicts that NAD+, FAD+ and electron transport all to be impaired (**Figure 4A**). Since Complex I is a major source of cellular ROS (32), we assessed ROS levels generated by WT and *as3mt*^*-/-*^ mutants at 120 hpf and found them decreased by an average of 38% in mutants (**Figure 4B**). Impaired mitochondrial function is a primary cause of non-alcoholic fatty liver disease (NAFLD) (33-35). We found that 81% of *as3mt*^*-/-*^ mutants developed fatty liver (steatosis) by 120 hpf (Figure 5C). Steatosis incidence and severity were fully rescued in *as3mt*^-/-^ mutants when As3mt expression was reintroduced only in hepatocytes by crossing to Tg-*as3mt*^*wt/tg*^ (**Figure 4F-H**). Thus, the transcriptional effects of *as3mt* mutation translates to functional consequences for the liver, including causing fatty liver. Together, these data unequivocally demonstrate that As3mt has an arsenic-independent role in the liver.

**Figure 5.**
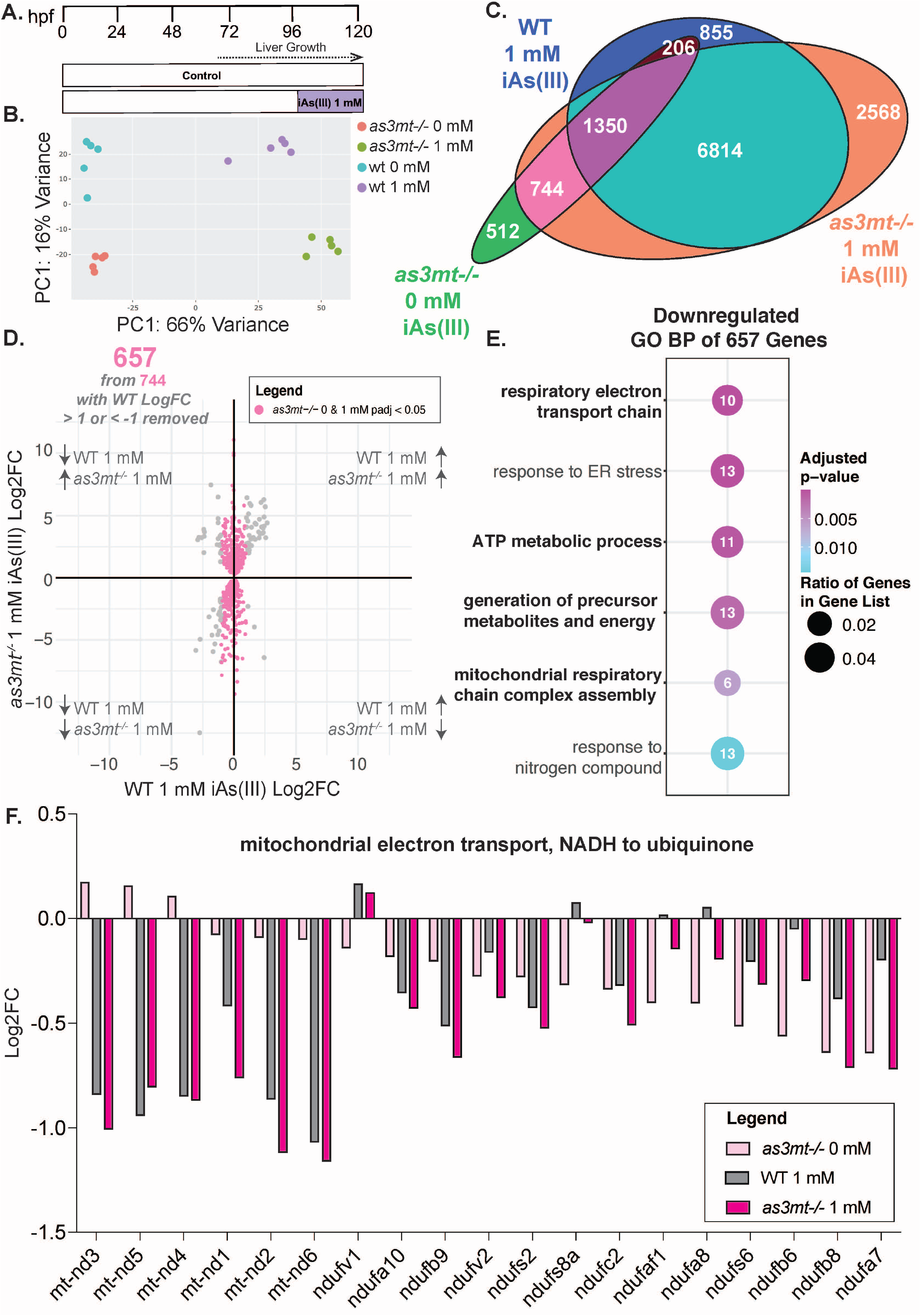
Loss of *as3mt* induced mitochondrial dysfunction dictates greater cellular sensitivity to iAs(III) rather than iAs(III) accumulation during acute iAs exposure. **A**. Treatment scheme of 1 mM iAs(III) from 96-120 hpf. **B**. PCA plot of all samples. **C**. Overlap of DEGs from untreated *as3mt*^*-/-*^, 1 mM treated WT, and 1 mM treated *as3mt*^*-/-*^. **D**. Cross plot of DEGs unique to untreated and 1 mM treated *as3mt*^*-/-*^. **E**. GO BP of pink genes from D. **F**. Log2FC of genes from *mitochondrial electron transport, NADH to ubiquinone*.

### Loss of *as3mt* sensitizes to iAs(III) by deregulating mitochondrial function

While AS3MT metabolism of arsenic is critical for reducing iAs(III) accumulation, we hypothesized that loss of *as3mt* can also contribute to iAs(III) toxicity by abrogating mitochondrial function, a central cellular target of arsenic (36). We carried out RNAseq analysis of the livers from WT and *as3mt*^-/-^ mutant larvae treated with 1 mM iAs(III) from 96-120 hpf and normalized each sample to untreated larvae of the same genotype (**Figure 5A**;**Table S6-7**). While there were significant differences based on both treatment and genotype (**Figure 5AB**), 88% of iAs(III) induced DEGs in WT livers were also differentially expressed and strongly linearly correlated in *as3mt*^-/-^ mutant livers (r = 0.937; **Figure 5C and S6A**). These genes were enriched in multiple pathways, including mitochondrial processes such as fatty acid beta-oxidation and RNA processing (**Figure S6B**). This indicates that the massive transcriptional response to iAs exposure is caused by the effects of iAs(III) on cellular homeostasis, and that the process of iAs(III) metabolism has little effect on the gene expression changes observed.

We reasoned that the increase in arsenic toxicity in *as3mt*^-/-^mutants was attributed either to a process that occurred in WT livers during iAs(III) exposure but was hindered in mutants, or to a process that occurred only in the mutants. We investigated this by analyzing the As3mt dependent DEGs – i.e those 882 genes which had no change (log2 fold change < -1 or > 1) in expression in *as3mt*^*-/-*^ mutants treated with iAs(III) but were significantly changed in WT treated samples. Surprisingly, GO analysis did not find any cellular pathway or cellular component that were enriched for this geneset. We next analyzed the 1,862 genes that had no change in expression (log2 fold change < -1 or > 1) in WT livers exposed to iAs(III) but were significantly changed in *as3mt*^-/-^ mutant livers only in response to iAs(III) (**Figure S6C**). These genes function in multiple unrelated pathways including ncRNA processing, transcription and glycosylation (**Figure S6D**). It is unclear to what extent these pathways are a response to iAs(III) accumulation or due to some other cellular defect caused by loss of As3mt.

We hypothesized that the disruption of mitochondrial function by As3mt deficiency (**Figures 2-3**) could impact the response to iAs(III) and, if so, this dependency would be reflected in the transcriptome of *as3mt*^*-/-*^ mutants exposed to iAs(III). We analyzed the 657 DEGs common to untreated and iAs(III) exposed *as3mt*^*-/-*^ mutants which showed no change in expression in WT larvae exposed to iAs(III) (**Figure 5D**). Notably, the downregulated genes were enriched in multiple processes relevant to mitochondrial function (**Figure 5E**). Pathway analysis of the unified set of DEGs in the liver of untreated and iAs exposed *as3mt*^*-/-*^ mutants plus WT iAs exposed larvae revealed that only two pathways were shared in all conditions: lipid metabolic process and mitochondrial electron transport, NADH to ubiquinone (**Figure 5E, Figure S6E**), but only the latter pathway was more affected in iAs(III) exposed *as3mt*^*-/-*^ mutants. Strikingly, 12 of the 19 nuclear and mitochondrially encoded genes that function in the NADH to ubiquinone pathway (i.e. Complex I) were downregulated in response iAs(III) exposure in WT larvae and were further downregulate in iAs(III) exposed *as3mt*^*-/-*^ mutants (**Figure 5F**). Thus, As3mt loss impairs mitochondrial function which further exacerbates the transcriptional response to iAs(III) exposure and, potentially, confers susceptibility to arsenic.

## DISCUSSION

To date, As3mt has been almost exclusively studied as a dedicated iAs(III) methyltransferase. The data presented here open new areas of investigation for understanding how arsenic causes disease and also identifies As3mt as a previously unrecognized regulator of hepatic energy metabolism, ROS homeostasis and contributor to fatty liver. We found that *as3mt*^*-/-*^ mutant zebrafish have thousands of DEGs in the whole animal which are even more profound in the liver, many of which persist to adulthood. *as3mt*^*-/-*^ mutants develop fatty liver, which makes this finding clinically relevant as close to a third of the world’s population is estimated to have fatty liver (37). Transcriptomic analysis uncovered a novel and essential role for As3mt in regulating mitochondrial function, specifically the electron transport chain, the TCA and ROS generation, with a prediction that oxidative phosphorylation and ATP generation are disrupted. This analysis also pointed to ribosome biogenesis and function as downregulated in *as3mt*^*-/-*^ mutants, but the functional significance of that finding is not yet clear, but it is interesting that neurons depleted of AS3MT also show deregulation of the same pathways (22). Overall, this study indicates that the high levels of As3mt detected in hepatocytes is important for maintaining hepatic metabolism in the absence of iAs.

These data challenge the current paradigm that As3mt solely functions in iAs(III) detoxification. Our findings are also directly relevant to understanding iAs toxicity, as iAs causes mitochondrial defects by targeting the same pathways that we discovered to be dependent on As3mt (36). Finally, we provide a novel perspective on the wealth of data from epidemiolocal and experimental studies which have shown that iAs is less toxic in people who have higher As3mt levels, and is more toxic when As3mt is reduced or deleted (38-40). We report that while *as3mt*^-/-^ mutants are sensitized to iAs(III) exposure, the majority of the gene expression changes caused by iAs are not different in *as3mt*^-/-^ mutants suggesting that most iAs responsive genes are largely due to iAs(III). We propose that impaired mitochondrial function caused by As3mt deficiency in developing larvae synergizes with the effects of iAs(III) on the mitochondria, exacerbating toxicity (Figure 8A). This study not only uncovers a new disease relevant function for a long-studied enzyme, but it also suggests a novel mechanism by which an endemic toxicant causes pathology.

Our working model is that As3mt regulates the expression and/or function of many of the proteins in Complexes I-IV, the TCA cycle and oxidative phosphorylation (Figure 8A). The downregulation of NADH-ubiquinone oxidoreductase *(ndufs)* genes that are essential for electron transport and function in Complex I in *as3mt* mutants predicts that coenzyme Q (CoQ) is oxidized, and unable to serve as the electron acceptor from Complex I. This is significant because CoQ has been implicated as a contributor to fatty liver in humans and animal studies (41), and CoQ supplements have been shown to protect against acetaminophen hepatotoxicity (42), lowered lipid levels in other models, and decreased signs of liver damage in patients with fatty liver (43). We propose a potential therapeutic application of CoQ for arsenic toxicity.

Our study provides a direct link between *as3mt* and fatty liver. It is well established that mitochondrial function is decreased in patients with fatty liver (44) and that mouse models of mitochondrial dysfunction and patients with mitochondrial disorders are characterized by fatty liver (45). We propose two mechanisms by which this happens (**Figure 6B**). First, fatty acid beta-oxidation is the main mechanism by which triglycerides are converted to ATP in hepatocytes. Thus, steatosis is a common outcome of toxicants or genetic perturbations that impair mitochondrial function. Second, downregulation of *pemt*, a key methyltransferase involved in phosphatidylcholine (PC) synthesis has been shown to reduce PC levels and resulting in steatosis in *pemt* KO mice (46). This is attributed to a failure of triglyceride packaging and export. In line with this, As3mt knock out mice have a complex pattern of changes to the levels of distinct PC species, suggesting that *As3mt* has uncharacterized roles in liver PC metabolism (18). We propose that downregulation of *pemt* in As3mt deficient cells both reduces PC synthesis, reducing VLDL packaging and lipid export (**Figure 6A**). Another study showed that AS3MT interacted with a component of the inflammasome and that this interaction contributed to steatosis following iAs exposure in mice (47), however we have not found this pathway to be disrupted in our studies. Further examination of the role of these pathways and of lipid flux in As3mt deficient hepatocytes will be important in future studies on the mechanism of steatosis in this model.

**Figure 6.**
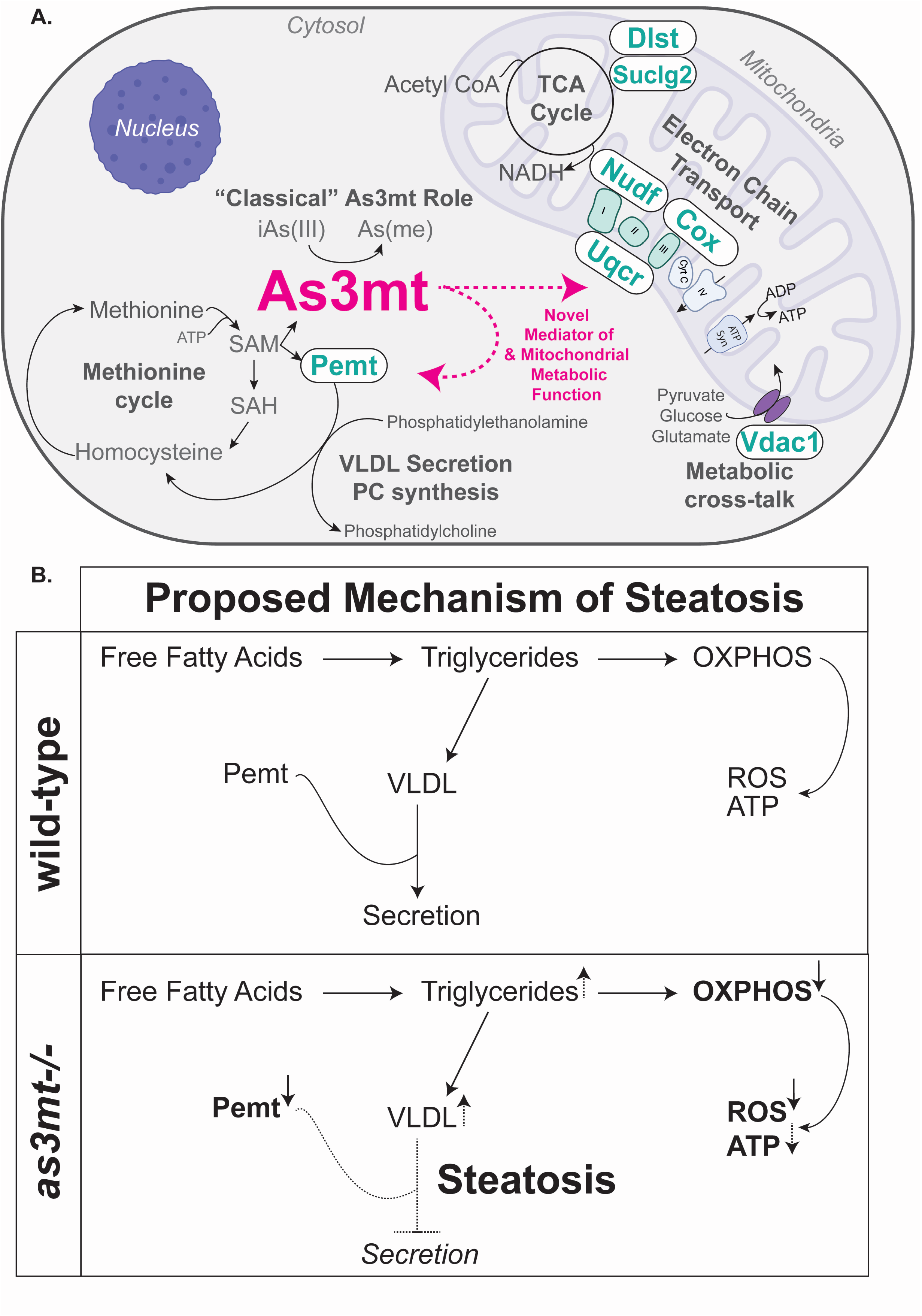
Proposed mechanism of loss of *as3mt* induced hepatic steatosis.

It is not clear what As3mt functions contribute to deregulation of mitochondrial metabolism and fatty liver. Given the highly conserved methyltransferase domain in As3mt, which retains homology to both protein and RNA methyltransferases, we speculate that As3mt directly modulates upstream regulators of proteins in the mitochondrial membrane machinery through methylation. One study implicated an RNA methyltransferase in the hepatic response to arsenic (47), but it is not clear whether AS3MT also functioned in this methylation reaction. Indeed, other methyltransferase activity in Complex I assembly and function have been demonstrated as essential for zebrafish and knockdown of these proteins results in more severe larval phenotypes, specifically in delayed hatching and overall morphology (48). Many mitochondrial proteins are regulated by methylation (49), and it is possible that As3mt contributes to these modifications. Alternatively, *as3mt* may impose an indirect effect in mitochondrial function by modulating the metabolites that feed into the electron transport chain. In line with this hypothesis, we observed downregulation *vdac1*, which aids in exchange of key components including nucleotides, pyruvate, malate, succinate, NADH/NAD+, as well as lipids, cholesterol and ions. Vdac1 acts as a major hub for crosstalk between metabolism, apoptosis, signal transduction, and anti-oxidation protein and, in a mouse model of fatty liver, supplementing Vdac1 improves beta-oxidation, reduces steatosis associated hepatic pathology (50). Which specific metabolic regulators are required for the effects observed in As3mt deficient hepatocytes requires further investigation.

Surprisingly, *as3mt* was transcriptionally downregulated in the liver during iAs(III) exposure. This would predict that iAs methylation would be decreased as exposure continues. Moreover, as iAs(III) accumulates, As3mt is diverted from its cellular function, and that further downregulation of *as3mt* expression could exacerbate the cellular defects caused by loss of As3mt function. We predict this to have significant consequences for hepatic toxicity as both the damaging effects of iAs(III) accumulation combined with the loss of As3mt functions are relevant. Moreover, this finding provides a potential mechanism for how iAs exposure contributes to fatty liver (11).

Finally, the findings presented here have major implications for iAs(III) endemic populations. To date, protective and maladaptive SNPs in the *AS3MT* locus in humans have been studied only in the context of iAs(III) metabolism, using serum or urinary iAs(III) and methylated species as an endpoint. Here, we find that *as3mt* has physiological roles which alone can augment iAs toxicity and our data showing that the transcriptomic response to iAs is largely similar in the presence or absence of As3mt suggest that loss of the arsenic metabolism function has less of an effect than its cellular, arsenic independent function. Future studies into *as3mt* expression and mitochondrial dynamics and downstream metabolic outputs will greatly aid the field of iAs toxicology and opens avenues for new iAs therapeutic strategies.

## MATERIALS & METHODS

### Zebrafish husbandry, embryo rearing and iAs exposure

All procedures were approved by and performed in accordance with the New York University Abu Dhabi Institutional Animal Care and Use Committee guidelines. Adult zebrafish were maintained on a 14:10 light:dark cycle at 28°C. Embryos were collected from natural spawning of group mating within 2 hours and were reared at 28°C. Experiments were conducted in 6-well plates (Corning, USA) with 20 embryos in 10 mL embryo medium, as described (24). Embryos were exposed to sodium meta-arsenite (Sigma-Aldrich, 7784-46-5; iAs (III)) by diluting 0.05 M stock solution to final concentrations ranging from 0.5 mM – 1.5 mM in embryo medium from either 6-120 hpf or 96-120 hpf as described (24). For all experiments, mortality was scored daily, dead embryos and larvae were removed upon identification, and morphology was recorded at 120 hpf as readout of iAs toxicity. After 120 hpf, all treated larvae were washed 3 times in fresh embryo media prior to downstream investigations.

A note on nomenclature: As a default, we use human nomenclature for genes and proteins and use species relevant nomenclature for statements that refer to data from a specific organism.

### as3mt^-/-^ Mutant Generation

sgRNA and primers targeting the *as3mt* gene was designed using ChopChop (https://chopchop.cbu.uib.no/) (**Table S8**). sgRNA was produced using sgRNA IVT kit (Takara Bio) by following the manufacturer’s instructions and RNA was isolated by Trizol (Invitrogen). sgRNA was quantified by Qubit RNA BR kit and diluted at 50 ng/*μ*l and stored as single use aliquots. WT-ABNYU embryos were injected with 1 nl of equal volume of previously diluted nls-Cas9 protein (IDT; 0.5 *μ*l of nl-Cas9 added with 9.5 ul of 20 mM HEPES; 150 mM KCI, pH 7.5) and sgRNA (incubated at 37° C for 5 minutes). At 96 hpf, 20 embryos were individually collected and genomic DNA was extracted by heat shock denaturation in 50 mM NaOH (95°C for 20 minutes). For each embryo, PCR was performed on genomic DNA by using Q5 High-Fidelity Taq Polymerase (New England Biolabs) followed by T7 endonuclease I assay (New England Biolabs) assay to detect mutations. For T7 endonuclease I assay, 10 *μ*l of PCR product was incubated with 0.5 *μ*l of T7e1 enzyme (New England Biolabs) for 30 minutes at 37°C. When the embryos reached sexual maturity (around 3 months), individual adult (F0) zebrafish were crossed to WT adults to generate the F1. DNA from putative was used to perform Sanger sequencing. Allele B F0 had an 8 bp indel mutation that was predicted to be a frameshift mutation. F1 embryos carrying this allele were used to generate *as3mt*^*-/-*^ mutants.

### Tg-as3mt^wt/tg^ Transgenic Generation

Hepatic overexpression of *as3mt* in transgenic zebrafish *(Tg(fabp10a:zas3mt; cry:dsRed))* were generated by injecting a vector that contains 2,813 bp of the liver promoter, *fabp10a*, upstream of upstream the coding sequence of zebrafish *as3mt* as determined from alignment with the zebrafish reference genome. The transgene cassette was flanked by *tol2* sites and the vector was injected into 1-2 cell stage embryos along with transposase mRNA. Larvae positive for red lens were selected and raised to adulthood and outcrossed to WT (TAB 5) adults to generate the line.

### Sanger Sequencing

Genomic DNA was extracted from individual embryos by heat shock denaturation in 50 mM NaOH (95°C for 20 minutes) and used for PCR by using Q5 High-Fidelity Taq Polymerase (New England Biolabs). The presence of specific amplicons was tested by running the PCR product on 1% agarose gel. 5 *μ*l of PCR products were purified by ExoSAP-IT™ PCR Product Cleanup Reagent (Thermo Fisher Scientific) following manufacturer’s instruction. Purified PCR products were sequenced using Sanger Sequencing Kit (Applied Biosystem) following manufacturer’s instruction and loaded on SeqStudio Genetic Analyzer (Applied Biosystems). Results were visualized on SnapGene Viewer to assess the quality of the run and analyzed in Synthego (https://ice.synthego.com/#/) to identify mutants.

### Gene Expression Analysis

Pools of at least 5 livers were microdissected from 120 hpf zebrafish larvae with transgenic marked livers (*Tg(fabp10a:Caax-eGFP))*. Larvae were anesthetized in tricaine and immobilized in 3% methyl cellulose and the livers were removed using 30-gauge needles. RNA was extracted from livers using TRIzol (Thermo Fisher, 15596026) and precipitated with isopropanol. For RNAseq analyses, all samples were DnaseI (ThermoFisher) treated. For library preparation, high quality of total RNA was QCed using Bioanalyzer (Agilent 2100; Agilent Technologies, Santa Clara, CA, USA) and quantified by Qubit fluorometer. Only RNA with RIN score >7 was used for library preparation. mRNA library was prepared using Illumina TruSeq V2 RNA sample Prep Kit (San Diego, CA) according to the manufacturer’s protocol. Briefly, 100 ng of total mRNA was poly-A purified, fragmented, and first-strand cDNA reverse transcribed using random primers. Following second-strand cDNA synthesis, end repair, addition of a single A base, TrueSeq adapter-index was ligated to cDNA libraries, and PCR amplification of 12 cycles was done for enrichment, producing a 350-400 bp fragment including adapters. The fragment sizes and purity of the libraries were confirmed by Bioanalyzer 2100 (Agilent Technologies). The quantities of the libraries required for RNAseq were determined by real-time qPCR using a KAPA library quantification kit. Enriched cDNA libraries were sequenced using the Illumina NextSeq550 (Illumina).

Raw FASTQ were first assessed for quality using FastQC v0.11.5. The reads where then passed through Trimmomatic v0.36 for quality trimming and adapter sequence removal, with the parameters (*ILLUMINACLIP: trimmomatic_adapter*.*fa:2:30:10 TRAILING:3 LEADING:3 SLIDINGWINDOW:4:15 MINLEN:36*). The trimmed read pairs were then processed with Fastp to remove poly-G tails and Novaseq/Nextseq specific artefacts. Following the quality trimming, the reads were assessed again using FastQC.

Post QC and QT, the reads were aligned to the *Danio rerio* reference genome GRcZ10 / ENSEMBL release 84 using HISAT2 with the default parameters and additionally by providing the *–dta* flag. The resulting SAM alignments were then converted to BAM format and coordinate sorted using SAMtools v1.3.1. The sorted alignment files were then passed through HTSeq-count v0.6.1p1 using the following options (*-s no -t exon -I gene_id*) for raw count generation. Concurrently, the sorted alignments were processed through Stringtie v1.3.0 for transcriptome quantification. Briefly the process looks like this, stringtie -> stringtie merge (to create a merged transcriptome GTF file of all the samples) -> stringtie (this time using the GTF generated by the previous merging step). Finally, Qualimap v2.2.2 was used to generate RNAseq specific QC metrics per sample.

For quantitative reverse transcription PCR (qRT-PCR), RNA was reverse transcribed with qScript (QuantaBio, 95048-025) and performed using Maxima Sybr Green/ROX qPCR Master Mix Super Mix (Thermo Fisher, K0221). Samples were run in triplicate on QuantStudio 5 (Thermo Fisher). Target gene expression was normalized to *ribosomal protein large P0* (*rplp0*) using the comparative threshold cycle (ΔCt) method. Primers for the genes of interest are listed in **Table S9**. Expression in treated or genetically altered animals was compared to untreated and/or WT controls to determine fold change. All datasets are publicly available on GEO (GSE228754 and GSE156419).

### Image Acquisition

For whole mount imaging of live larvae, embryos were anesthetized with 500 *μ*M tricaine (Ethyl 3-aminobenzoate methanesulfonate; Sigma-Aldrich), mounted in 3% methyl-cellulose on a glass slide and imaged on a Nikon SMZ25 stereomicroscope.

Nile Red powder (Invitrogen) was dissolved in methanol at a concentration of 1 mg/*μ*L as the stock solution. At ∼119 hpf, larvae were treated with 20 *μ*L of Nile Red for 1 hour. Fixed imaging was performed after the larvae were fixed at 120 hpf in 4% paraformaldehyde (UTECH). Before imaging, the larvae are washed with PBS (UTECH). Fixed larvae were transferred to 0.17 mm imaging plates (FlouroDish) and embedded in 1% low melting agarose gel (SeaPlaque Agarose, Lonza) and imaged using a super resolution microscope-Leica STED 3X at 63x water lens.

### Steatosis Analysis

The incidence and severity of steatosis in livers was determined using Imaris. The Area for each liver section was manually outlined and the number of lipids droplets was determined using the spot count function in Imaris. The number of spots was divided by the surface area of individual livers. To obtain the incidence, the percentage of livers with number of spots > 2 was divided by the total number of livers imaged per clutch.

### IC-ICP-MS

At 120 hpf, after 6-120 hpf iAs(III) exposure, 20 live larvae from each condition were collected and washed 5 times with embryos water, with all excess liquid removed. Tubes were immediately freeze dried (Lypholizer - Christ Alpha 1-2 LD plus) and stored at -80. Prior to sonication, 500 µL of mobile phase (10 mM NH_4_H_2_PO_4_ (Sigma, 7722-76-1) in 5% Methanol, pH 7.9.) was added to each tube. Samples were sonicated with a probe sonicator (Branson) (2 min, 01 on, 02 off, 30% amplitude) on ice. Finally, samples were filtered (0.2 uM; Pall Corp; 4552T) and quantified within 24 hours of preparation.

All measurements were carried out with an Agilent 7800 ICP-MS instrument (ICP-MS/Agilent Technologies, Japan), equipped with a MicroMist nebulizer and a Peltier-cooled (2 ∘C) scott-type spray chamber for sample introduction. For arsenic speciation studies, the metrohm 940 Professional IC Vario, an anion exchange column (RP-X100, 250 mm × 4.1 mmi.d., Hamilton, USA) were used for anion exchange column liquid chromatography. Softwares used are MagicNet IC to interface with the ion chromatography system, masshunter for ICP-MS (with chromagraphic analysis and advanced acquisition activated) and Maestranova with MS plugins for data analysis. Ultrapure water (18MΩ cm resistivity) was obtained from an integral 10 miliq water purification system (Millipore, Bedford, MA, USA). Standard 4 element, Agilent ICP-MS tuning stock solution was used for tuning and calibration. For Quantitative determination, solutions were prepared from ICP-MS standard stock for As at 1000 ppm (Agilent). IC-ICP-MS standards were: As(III) (Arsenic (III) Standard, Fisher Scientific; 1327-53-3), As(V) (Arsenic V Speciation Standard, Fisher Scientfic; 7732-18-5, AsB (Arsenobetaine Standard Solution, LGC Standards; NIST-3033), DMA (Dimethylarsinic Acid Standard Solution, LGC Standards; NIST-3031) and MMA (Monomethylarsonic Acid Standard Solution, LGC Standards; NIST-3030). Using the 100 μL injection loop, samples were loaded onto the column (PRP-X100, 250 mm × 4.1 mm i.d., Hamilton) at a flow of 1 mL/min of mobile phase. The flow is analyzed using the ICP-MS with the following method conditions: Acquisition Mode (TRA), Plasma conditions: Radiofrequency power: 1550 W, tune mode: no gas, Plasma mode: General Purpose, Quick Scan: off, Independent, Nebulizer Gas: 1.23 L/min, Monitoring mass ^75^As. Analysis methods were run for 25 minutes.

### ROS Assay

ROS level changes were determined as previously described (12) with the use of 5-(and- 6)-chloromethyl-2′,7′-dichlorodihydrofluorescein diacetate, acetyl ester (CM-H_2_DCFDA; Invitrogen). Medium from larvae was placed in culture media containing 5 μM CM-H_2_DCFDA for 90 minutes. Fifty *μ*L of solutions were loaded onto 96 well plates in 3 technical replicates. The fluorescence of each well was determined using Synergy H1 Hybrid Multi-Mode Reader (BioTek), 485 nm and 530 nm emission (bottom read).

### Statistical analysis, rigor and reproducibility

All experiments were repeated on at least 2 clutches of embryos, when possible, with all replicates indicated. Reproducibility was assured by carrying phenotype scoring and other key experiments by independent investigators. Data are presented as normalized values. Statistical tests were used as appropriate to the specific analysis, including Student’s T-test, ANOVA and Chi Square (Fisher’s exact test) using Graphpad Prism Software or R to analyze the RNAseq data.

## Author Contributions

PD and KCS conceived the idea and planned the experiments. PD, NAK, MJO, EMAG, SP and KCS carried out the experiments and PD, KCS, NAK, MJO, EMAG, AS, SP and JT analyzed the data, PD prepared the figures and PD and KCS wrote the manuscript.

## Acknowledgements

The authors are grateful to Koichi Kawakami for the *(Tg(UAS:GFP;gSAlzGFFD886A))* zebrafish line, Soja Soman for the *as3mt*^*wt/tg*^ plasmid, Elena Magnani, Anjana Ramdas Nair, Eleanor Jenkins and Shashi Ranjan for technical support, and to all members of the Sadler lab for helpful discussions. The NYUAD Core Technology Platform Imaging and Genomics Facilities provided invaluable technical support. Gratitude for input from Kristin Gunsalus, Piergiorgio Percipalle, Andreas Hochwagen, and Max Costa.

## Supplemental Figures

**Figure S1. Endogenous *as3mt* is highly expressed and enzymatically active in zebrafish larvae. A**. Fluorescent images of endogenous *as3mt* transcription between 72-120 hours post fertilization (hpf). **B**. Overlay of brightfield and fluorescent images of a zebrafish larva at 120 hpf. **C**. Mean read counts from two RNAseq datasets of single livers from 120 hpf zebrafish larvae. *as3mt* is marked in red. **D**. IF of WT 120 hpf zebrafish liver with hoechst and anti-AS3MT. **E**. IC-ICP-MS data from 12.5, 3.125 and 0.75 ppb As standard mix (As(III), DMA, MMA, AsB and As(V)) and zebrafish extracts after exposure to 1 mM iAs from 6-120 hpf (black line).

**Figure S2. CRISPR/Cas9 mediated *as3mt* mutagenesis in zebrafish. A**. *as3mt* gene (16208 bp) with introns denoted as lines and exons as boxes. *as3mt* sgRNA targeted exon 3. **B**. 1% agarose gel of amplified *as3mt* exon 3 from 3 pools of 5 embryos (F1s) from CRISPR/Cas9 injected Founder **B**. PCR product was run without (-) and with (+) T7e1. **C**. 1% agarose gel of fin clips adults F1s from outcross of CRISPR/Cas9 injected Founder B without (-) and with (+) T7e1. Green denotes a positive carrier of edited *as3mt* whereas red denotes a negative carrier (WT) fish. **D**. 1% agarose gel of amplified *as3mt* from F2 fin clips with primers designed for either all of exon 3 (T7e1 primer), WT *as3mt*, or edited *as3mt*. Sanger sequencing results from a WT, heterozygous or homozygous *as3mt* edited gene above. **E**. Integrative Genomic Viewer (IGV) view of aligned RNAseq reads from F3 whole larvae from either an *as3mt*^*-/-*^ (top) or WT incross (bottom). RNA from *as3mt*^*-/-*^ larvae have an expected 8 bp deletion in exon 3 of *as3mt*. **D**. Translated RNA reads from WT or *as3mt*^*-/-*^ (MUT) larvae.

**Figure S3. A**. Cross plot of overlapped DEGS in *as3mt*^*-/-*^ whole larvae (x) and larval livers (y). **B**. GO BP analysis of blue genes from A.

**Figure S4. A**. PCA plot of male WT (purple) and *as3mt*^*-/-*^ (green) and female WT (blue) and *as3mt*^*-/-*^ (red) livers. B. Crossplot of DEGs from *as3mt*^*-/-*^ males (burnt orange) and females (teal). **C**. Venn overlay of DEGs in *as3mt*^*-/-*^ livers from larvae (purple), adult females (blue), and adult males (orange). **D**. Heatmap of 136 positively correlated genes from 85 + 50 + 162 overlay in D in larvae, males and female livers. **E**. Cross plot of shared DEGs in *as3mt*^*-/-*^ livers from males (x) and larvae (y). **F**. Raw count data from *pemt* and *vdac1*.

**Figure S5. A**. Representative images of hepatocytes at 120 hpf expressing nls-mCherry *(tg(fabp10a:nls-mcherry)*) in control (left) and *as3mt*^*wt/tg*^ larvae (rigtht). **B**. Quantification of number of nuclei per liver size, n = ∼15, 2 clutches. ns = not significant, unpaired t-test. **C**. Representative images of hepatocytes at 120 hpf expressing CAAX-eGFP *(tg(fabp10a:CAAX-eGFP))* in control (left) and *as3mt*^*wt/tg*^ larvae (right). **D**. Quantification of the area of hepatocytes, n = ∼12, 2 clutches. ns = not significant, unpaired t-test. **E**. Representative images of bile ducts *(tg(tp1:β-globin-eGFP))* in control (left) and *as3mt*^*wt/tg*^ larvae (right). **F**. Number of nodes from bile duct images n = 14-22, 2 clutches. **G**. Representative images of bile ducts *(tg(tp1:β-globin-eGFP))* and vasculature *(tg(flk1:ras-mCherry))* in control (left) and *as3mt*^*wt/tg*^ larvae (right). **H**. Represenative fluorescent image of 120 hpf control and *as3mt*^*wt/tg*^ larvae fed with the BODIPY-tagged phospholipid PED6. Quantification of positive gallbladder PED6 accumulation below (n = 12-15, 1 clutch).

**Figure S6. A**. Cross plot of shared DEGs in WT (x) and *as3mt-/-* (y) larval livers. **B**. - Log(FDR) of top 20 GO BP pathways in 1 mM iAs (III) treated WT (white) and *as3mt-/-* (pink) larval livers. **C**. Cross plot of DEGs unique in 1 mM iAs(III) *as3mt-/-* (y) sample compared to WT (x). **D**. GO BP analysis of genes from C. **E**. -Log(FDR) of GO BP analysis of shared pathways in untreated *as3mt-/-* (light pink) or 1 mM iAs (III) exposed WT (grey) and *as3mt-/-* (pink) samples.

