## Supplementary figures and images for "Arsenite methyltransferase 3 regulates hepatic energy metabolism which dictates the hepatic response to arsenic exposure"

### Figure S1

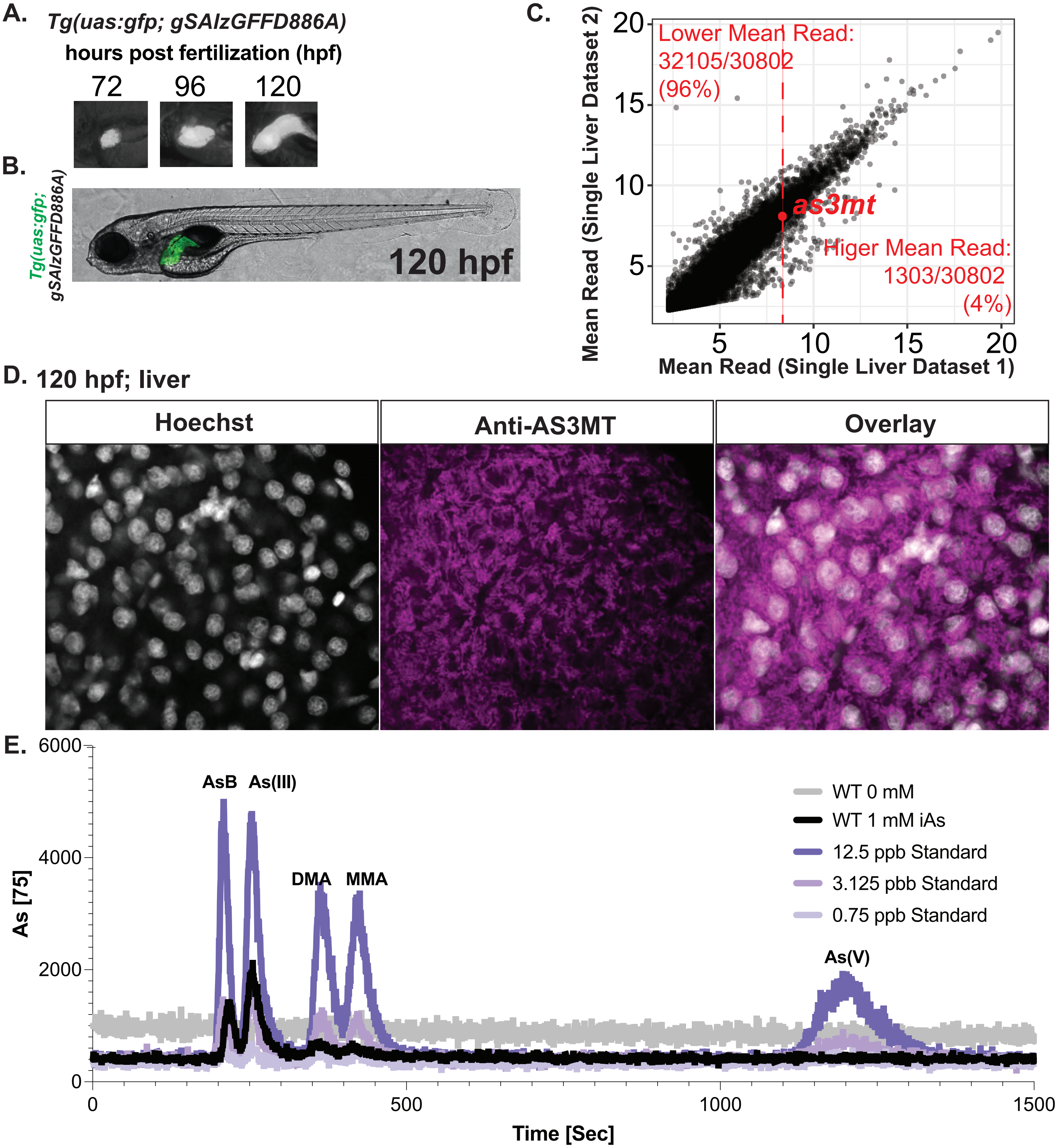

### Figure S2

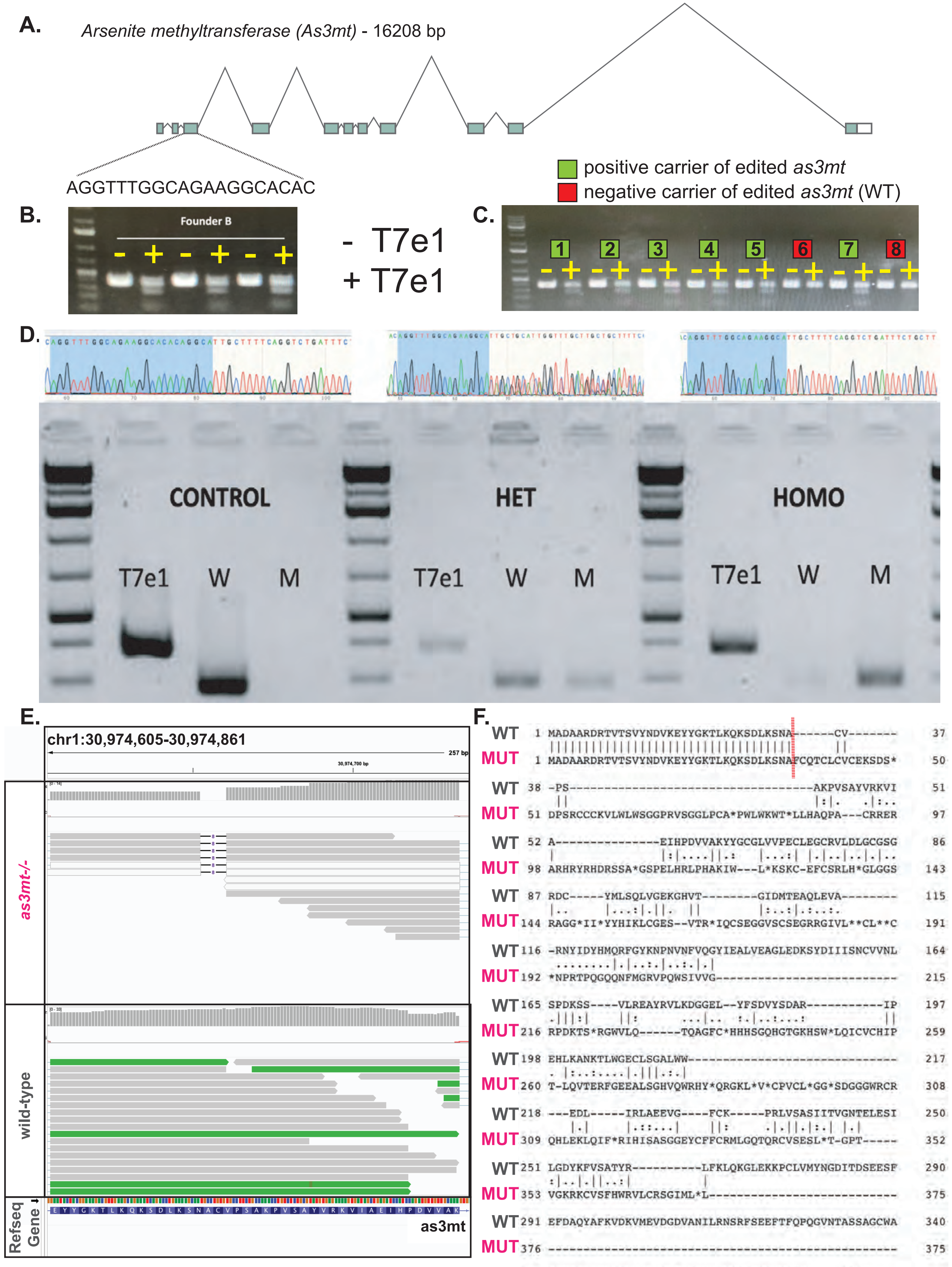

### Figure S3

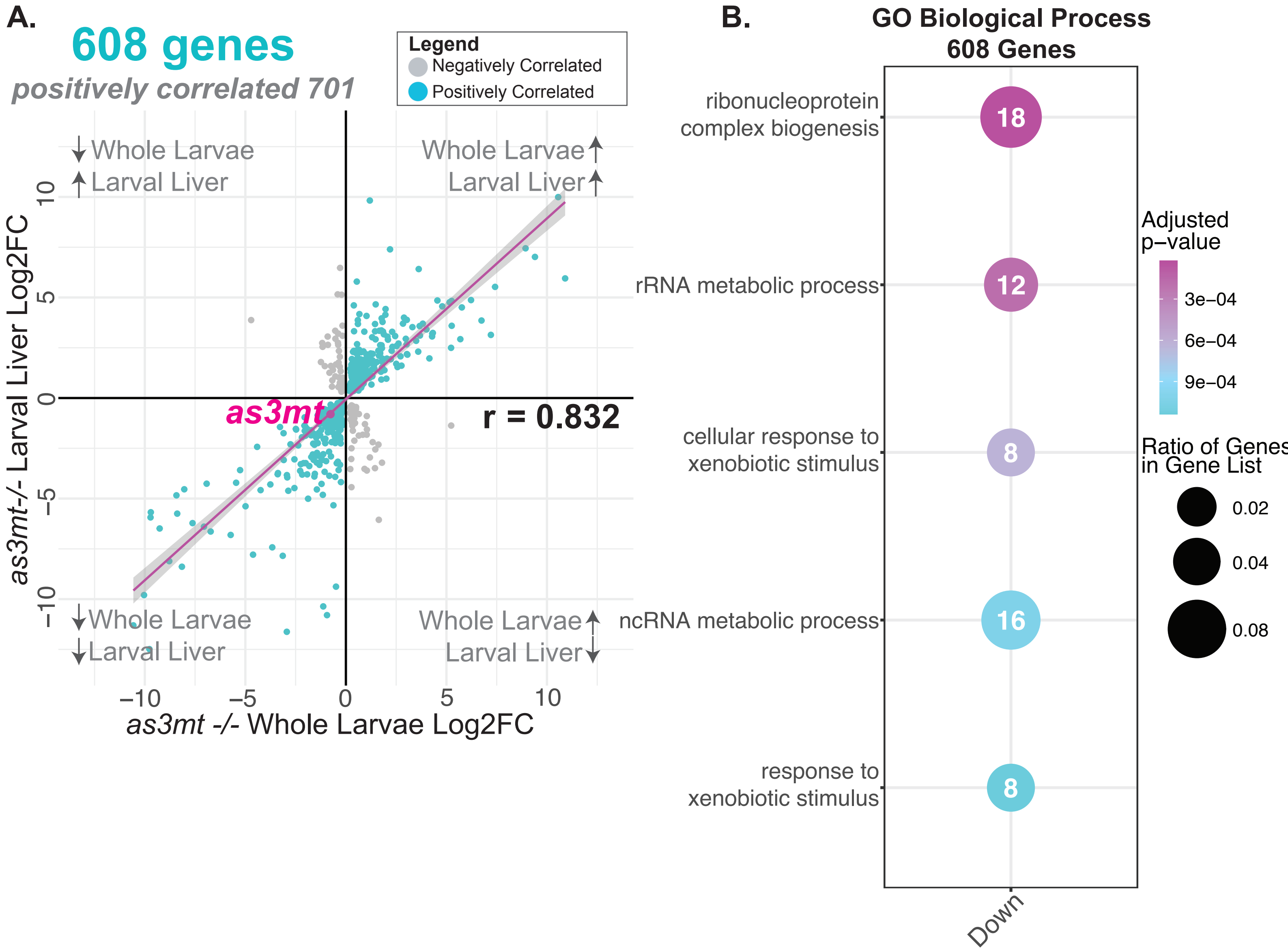

### Figure S4

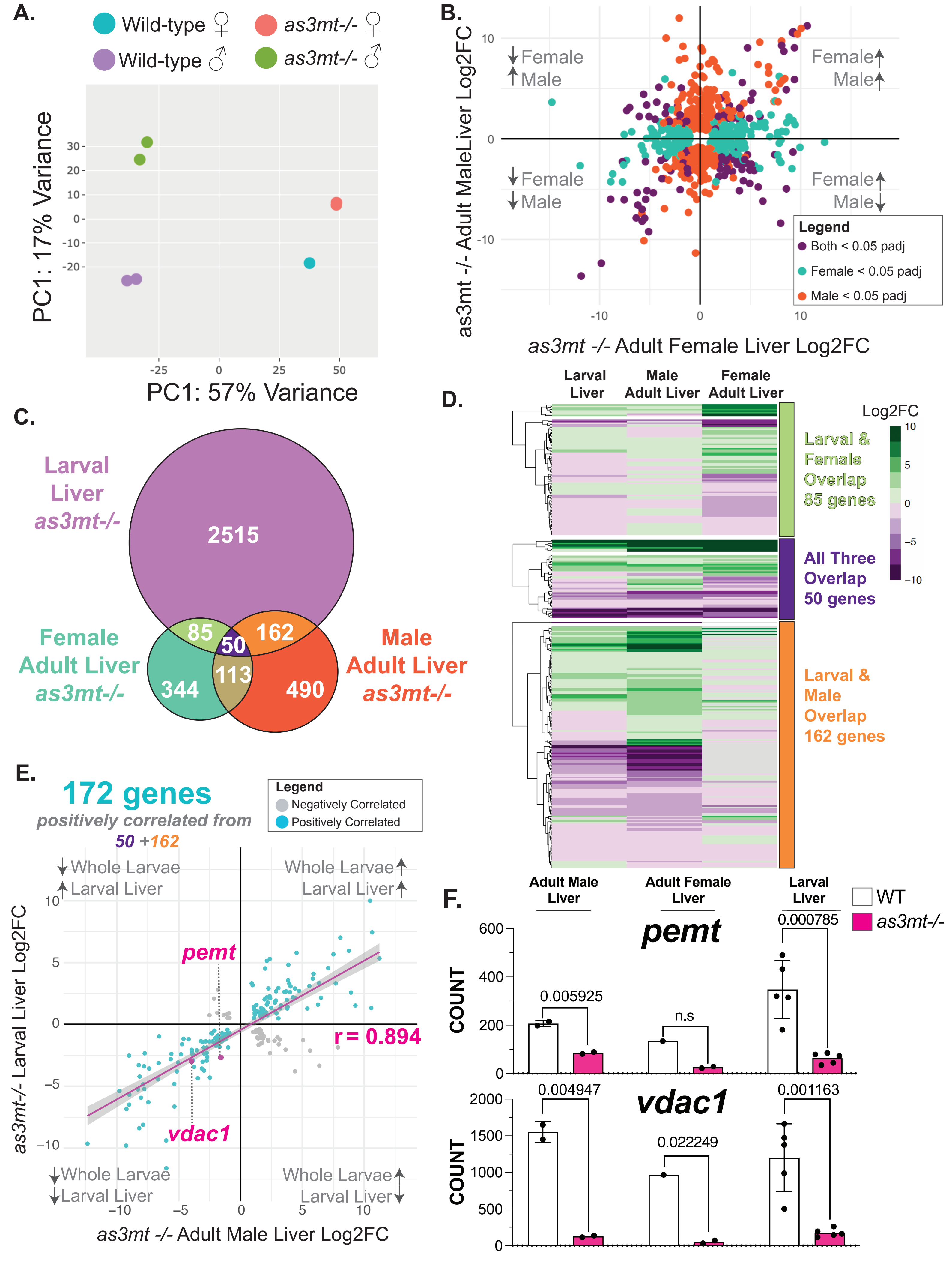

### Figure S5

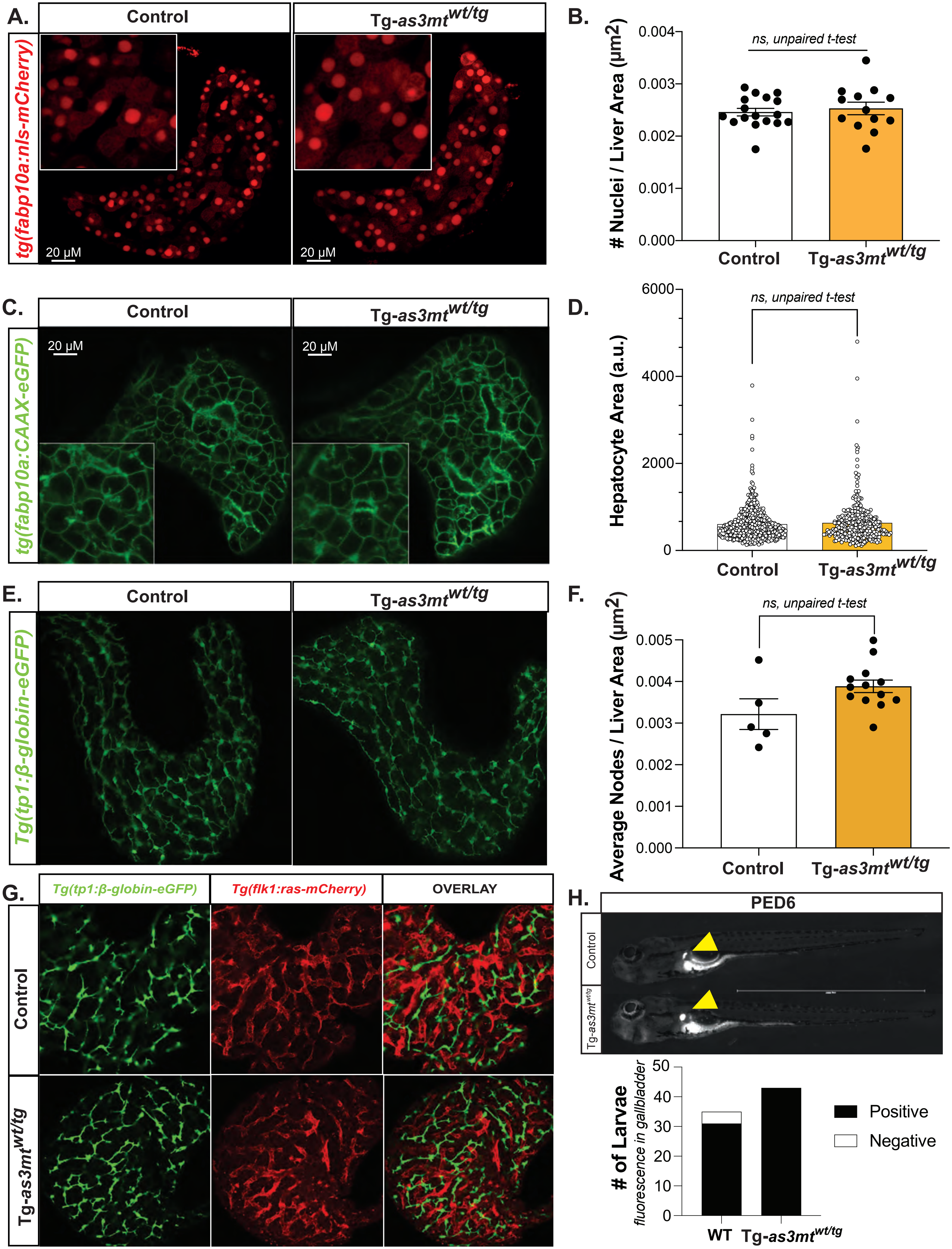

### Figure S6

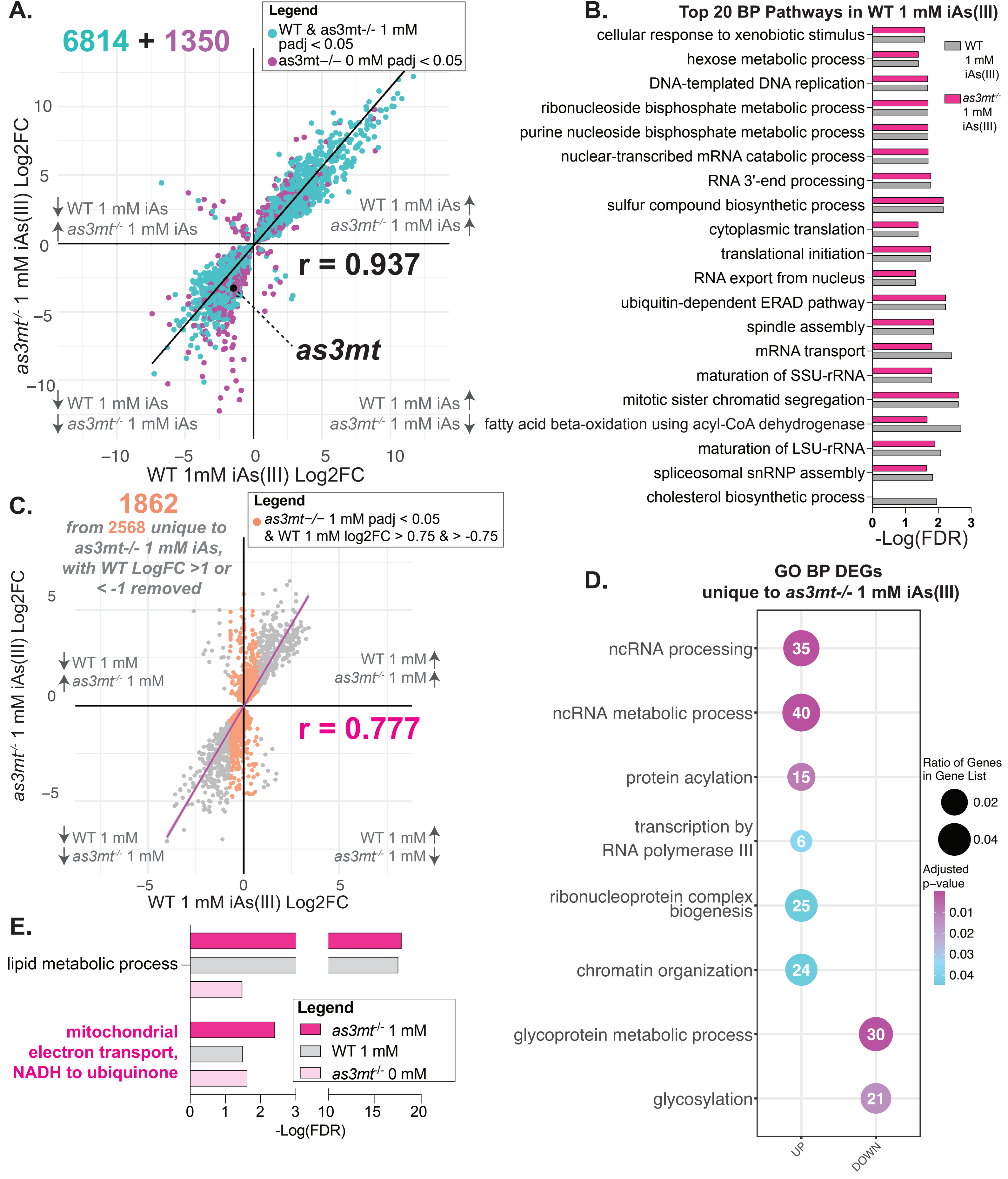
