## Supplementary material for "Arsenite methyltransferase 3 regulates hepatic energy metabolism which dictates the hepatic response to arsenic exposure": Table S8

| **Purpose** | **Forward 5’-3’** | **Reverse 5’-3’** |
| --- | --- | --- |
| sgRNA guide | GCGTAATACGACTCACTATAGgAGGTTTGGCAGAAGGCACACGTTTTAGAGCTAGAAATAGCAAGTTAAAATAAGGCTAGTCCGTTATCAACTTGAAAAAGTGGCACCGAGTCGGTGCTTT | AAAGCACCGACTCGGTGCCACTTTTTCAAGTTGATAACGGACTAGCCTTATTTTAACTTGCTATTTCTAGCTCTAAAACGTGTGCCTTCTGCCAAACCTcCTATAGTGAGTCGTATTACGC |
| T7e & SangerSeq | TCTTGTTCAAAGGGACAGGACT | ACCACATCAACACAACACATCA |
| WT genotyping | TCTTGTTCAAAGGGACAGGACT | GCAGAAGGCACACAGGC |
| MUT genotyping | TCTTGTTCAAAGGGACAGGACT | AGGTTTGGCAGAAGGCATTG |
| **Table S8. CRISPR guides and primers for *as3mt*** | | |
