## Supplementary material for "Arsenite methyltransferase 3 regulates hepatic energy metabolism which dictates the hepatic response to arsenic exposure": Table S9

| Ensembl ID | Symbol | Forward 5’-3’ | Reverse 5’-3’ |
| --- | --- | --- | --- |
| ENSDARG00000051783 | *rplp0* | CTGAACATCTCGCCCTTCTC | TAGCCGATCTGCAGACACAC |
| ENSDARG00000027572 | *as3mt* | CCGGGCTGGAGGATAAATCA | CCCTCCGTCCTTCAGAACAC |
| **Table S9. qPCR Primers** | | | |
